# Delta Oscillations Are a Robust Biomarker of Dopamine Depletion Severity and Motor Dysfunction in Awake Mice

**DOI:** 10.1101/2020.02.09.941161

**Authors:** Timothy C. Whalen, Amanda M. Willard, Jonathan E. Rubin, Aryn H. Gittis

## Abstract

Delta oscillations (0.5–4 Hz) are a robust but often overlooked feature of basal ganglia pathophysiology in Parkinson’s disease and their relationship to parkinsonian akinesia has not been investigated. Here, we establish a novel approach to detect spike oscillations embedded in noise to provide the first study of delta oscillations in awake, dopamine depleted mice. We find that approximately half of neurons in the substantia nigra reticulata exhibit delta oscillations in dopamine depletion and that these oscillations are a strong indicator of dopamine loss and akinesia, outperforming measures such as changes in firing rate, irregularity, bursting and synchrony. We further establish that these oscillations are caused by the loss of D2 receptor activation and do not require motor cortex, contrary to previous findings in anesthetized animals. These results give insight into how dopamine loss leads to dysfunction and suggest a reappraisal of delta oscillations as a biomarker in Parkinson’s disease.

## Introduction

Parkinson’s disease (PD) is characterized by the loss of dopamine neurons in the substantia nigra pars compacta (SNc), inducing a state of dopamine depletion (DD) in the basal ganglia. In human PD patients, this change is accompanied by a striking increase of oscillatory power in local field potential (LFP) recordings and in the spiking of individual neurons, primarily in the beta (13-30 Hz) and delta/theta frequencies (1-7 Hz) (Lenz et al., 1988; Levy et al., 2002; Priori et al., 2004; Steigerwald et al., 2008; Du et al., 2018; Halje et al., 2019).

Of these, beta oscillations have been the primary focus of research. In PD studies, beta oscillations have been shown to correlate with symptom severity (Jenkinson & Brown, 2011) and tend to dissipate under treatments such as dopamine replacement therapy (Weinberger et al., 2006; Ray et al., 2008). Similar oscillations are observed in some animal models of PD – slightly higher in frequency (25-35 Hz) in 6-hydroxydopamine (6-OHDA) lesioned rats or lower (8-13 Hz) in 1-methyl-4-phenyl-1,2,3,6-tetrahydropyridine (MPTP)-treated monkeys. In these models, the link between beta oscillations and motor symptoms is less clear. Beta oscillations may arise later than symptoms (Mallet et al., 2008), do not consistently track symptom progression (Muralidharan et al., 2016) or reduction with treatments such as deep brain stimulation (DBS) (McConnell et al., 2012), and occur in both parkinsonian and healthy animals (Connolly et al., 2015). Attempts to artificially induce beta oscillations in these animals have also been insufficient to cause PD-like symptoms (Swan et al., 2019). Even in humans, DBS studies have shown conflicting results between the correlation of beta oscillations and motor symptoms – oscillations tend to weaken during stimulation (Kühn et al., 2006, 2008) but not in every case (Rossi et al., 2008) or may return before symptoms reemerge (Foffani et al., 2006).

In contrast, the lower frequency oscillations observed in human PD patients have been much less studied, despite occurring in a greater number of neurons than beta oscillations in some patients or in the absence of beta oscillations at all (Levy et al., 2002; Du et al., 2018; Zhuang et al., 2019). These delta oscillations are often termed “tremor frequency” oscillations due to their typical coherence with Parkinsonian tremor (Bergman et al., 1994), but they may also arise without such coherence (Hurtado et al., 1999). While delta oscillations have an unclear relationship to tremor, their relationship to other PD symptoms such as bradykinesia and rigidity has not been investigated.

This lack of attention is surprising, as slower oscillations have also been observed in animal models of PD. In monkeys, oscillations as low as 3-7 Hz have been observed (Raz et al., 2000; Heimer et al., 2006; McCairn & Turner, 2009), but these have mostly been viewed as an extension of the beta band. In anesthetized rodents, oscillations at even lower frequencies (0.5– 4 Hz) are most prevalent (Tseng, Kasanetz, Kargieman, Pazo, et al., 2001; Walters et al., 2007; Parr-Brownlie et al., 2009; Aristieta et al., 2016), but these have been mostly discounted as artifacts of anesthesia or artificial respiration (Ruskin et al., 2002). Indeed, delta oscillations in the striatum of 6-OHDA-lesioned rats were shown to have high coherence to anesthesia-induced slow waves in motor cortex (M1) (Tseng, Kasanetz, Kargieman, Riquelme, et al., 2001; Belluscio et al., 2003) and were weakened after cortical ablation (Magill et al., 2001), leading to the conclusion that they merely infiltrate the basal ganglia through M1 and are not relevant to the awake, behaving parkinsonian animal. Experiments investigating the presence of sub-beta band oscillations in awake, behaving animals have, to our knowledge, not been performed.

One factor limiting these investigations is the high levels of noise that contaminate low frequency signals, particularly during awake recordings. So-called “pink” or “flicker” noise is most prevalent at low frequencies and typically observed in LFP recordings but is also present in the spiking of individual neurons. This complication makes reliable detection of oscillations near or below 2 Hz difficult with current methods.

Here, we develop a method to reliably distinguish spike oscillations from noise and use this approach to characterize the oscillations in the substantia nigra pars reticulata (SNr) of dopamine depleted mice. We demonstrate that delta, not beta, oscillations are the primary oscillatory feature in SNr neurons after loss of dopamine, and that they correlate strongly with PD-like motor deficits. We show that, contrary to prior reports, delta oscillations in the SNr precede those in M1, and that M1 is not necessary for these oscillations to develop in the SNr. We also establish that a loss of D2 receptor activation is sufficient to immediately and reversibly generate both delta oscillations and PD-like akinesia in awake mice, suggesting a direct link between dopamine loss, delta oscillations, and parkinsonian symptoms. This work indicates that delta oscillations in basal ganglia neurons are a critical component of parkinsonian pathology in DD mice and suggests that DD mice may effectively model the low frequency oscillations seen in PD patients.

## Results

### Dopamine depleted mice exhibit 0.5–4 Hz spike oscillations in SNr units

We recorded single units from the substantia nigra pars reticulata (SNr) of awake, head-fixed mice (Figure 1a–b) that had been bilaterally dopamine depleted with 6-OHDA or saline. To investigate oscillations in the spiking activity of single units, we first examined spike trains and their autocorrelograms. In control animals, units typically fired in a regular, pacemaking pattern, indicated by a fast oscillation in their autocorrelograms which corresponded to the interspike interval of pacemaking and flattened within 20-100 ms (Figure 1c). In contrast, units in bilaterally dopamine depleted animals exhibited autocorrelograms that showed much slower oscillations between 0.5 and 4 Hz that remained autocorrelated for several seconds, visible in the raw spike trains as peaks and troughs or pauses in firing (Figure 1d). These slow oscillations were never observed in the autocorrelograms of units from control animals.

**Figure 1.**
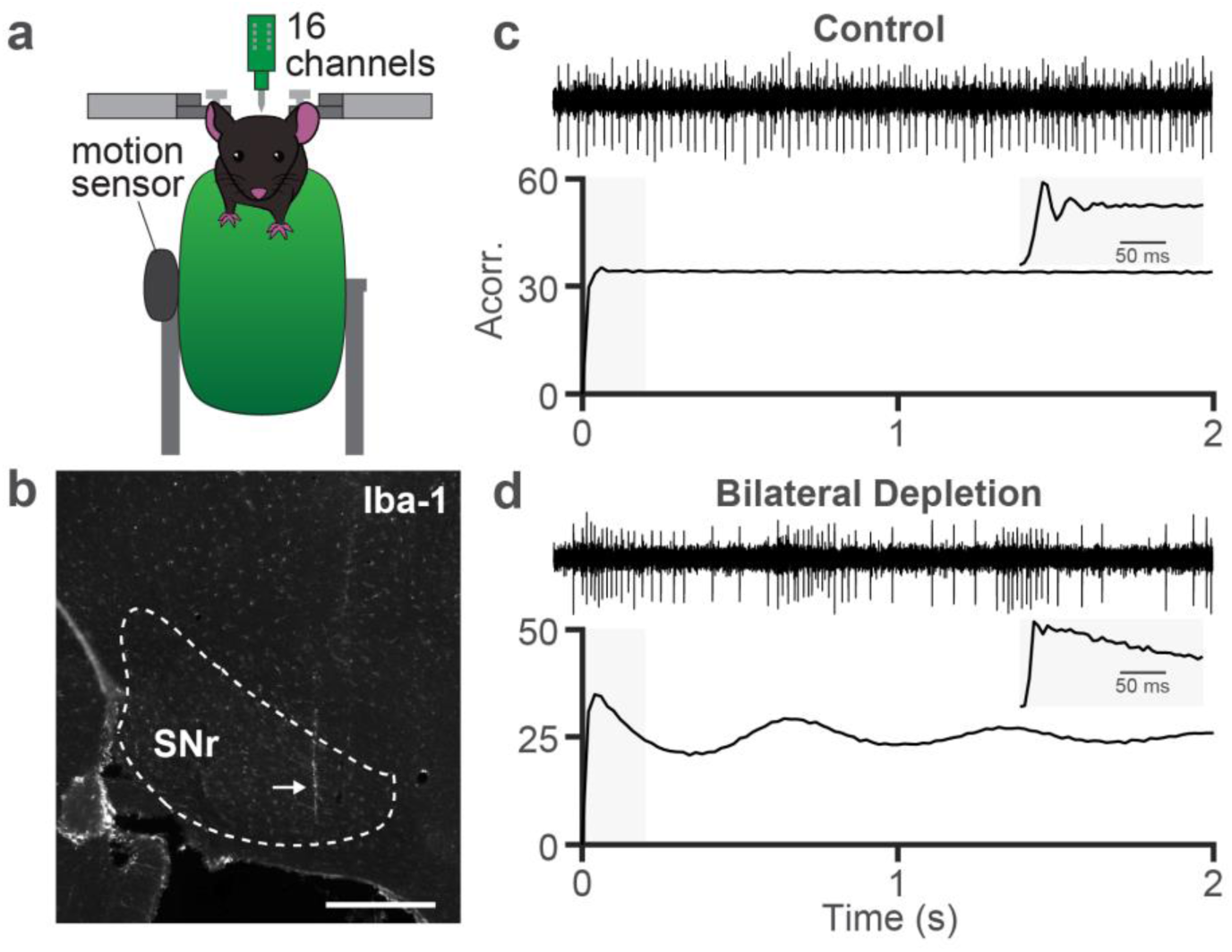
Dopamine depletion leads to low frequency spiking oscillations in SNr units. **a.** Schematic of recording setup. Mice were head-fixed atop a free-running wheel with attached movement sensor and single units were recorded with a 16-channel probe. **b.** Example sagittal slice with IBA immunofluorescence showing location of the recording probe in SNr. Dotted line indicates approximate location of target nucleus, arrow indicates probe location. Scale bar = 500 µm. **c.** Two seconds of an example SNr unit firing from a control animal (top) and the unit’s autocorrelation (bottom). Inset is zoomed into the first 200 milliseconds of the autocorrelation using a smaller bin size. **d.** Same as **c** for a bilaterally dopamine depleted animal.

### Phase shift analysis enables distinction between low frequency oscillations and neural noise

We first sought to reliably quantify these oscillations in dopamine depleted mice. Neural noise is more prevalent in awake than anesthetized animals, and typically manifests in a power law fashion (called “pink” or “flicker” noise) such that it is dominant in low frequencies. Since the oscillations we observe in SNr units in DD were in the range typically tainted by pink noise, we could not reliably detect them using standard approaches based solely on the power spectral density or transformations of it. Specifically, random peaks in the power spectral density atop pink noise, or the pink noise itself, can easily be misidentified as an oscillation of interest (Figure 2c).

**Figure 2.**
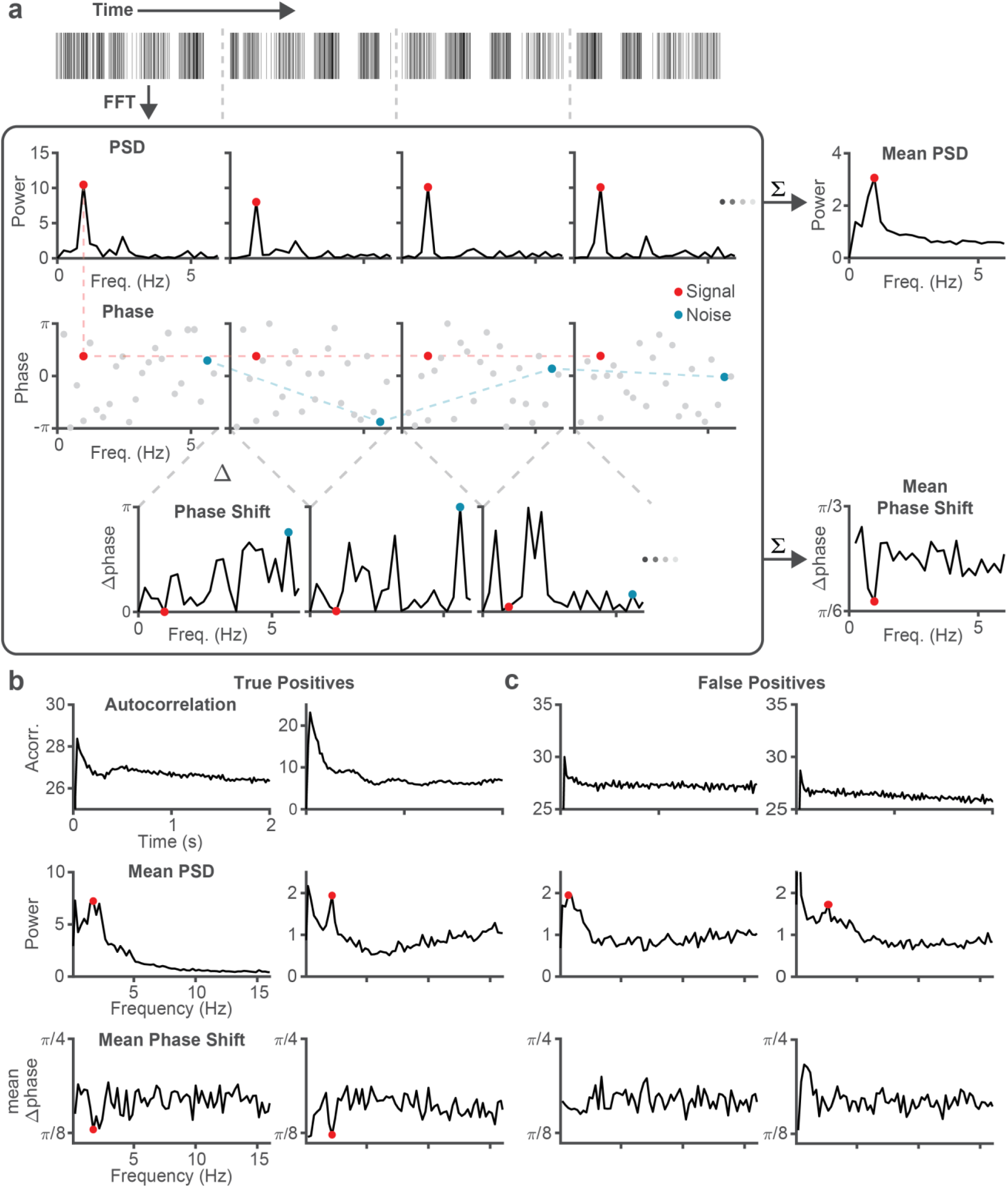
A phase shift measure to distinguish oscillations from noise. **a.** Diagram of the phase shift oscillation detection method. A spike train is divided into overlapping windows (1st row) and its Fourier transform is computed (corrected for the interspike interval distribution, see Methods). We identify statistically significant peaks in the 0.5-3 Hz range (compared to a control 100-500 Hz range) in the averaged power spectral density (PSD) across all windows (2nd row) and label the oscillation phase (3rd row) at that frequency. Notice while the peak frequency (red) has consistent phase across windows, an arbitrary noise frequency (blue) has inconsistent phase. We take the absolute circular difference of phases at each frequency (4th row) and compute whether the frequency identified in the power spectrum also has statistically significantly lower phase difference than the control band. A spike train which has both a significant spectral peak and significant phase difference trough at the same frequency is labeled as oscillating. **b.** Data from two example oscillating units. Top: Autocorrelation exhibiting oscillations. Middle: Significant peaks (red dots) in the PSD surrounded by pink noise. Bottom: The phase difference at these detected frequencies is significantly lower than control frequencies. **c.** Same as **b**, but for two units whose autocorrelation appears to be non-oscillating yet have a peak in their PSD and which would be “false positive” detections if only PSD’s were analyzed without the consideration of phase shift.

To overcome false positive detections, we used both the power and phase information provided by the short time Fourier transform to identify oscillatory components of spike trains with consistent phase over time (see Methods). By requiring that an oscillation have both high spectral power and low phase shift (Figure 2a), we successfully distinguished the oscillations of interest embedded in pink noise from the noise itself (Figure 2b). Notably, spike trains that exhibit a relatively flat autocorrelation but have delta peaks in their PSD are successfully disregarded as oscillators when phase shift analysis is applied (Figure 2c).

### Delta, not beta, oscillations in SNr units are a marker of dopamine depletion

Using this detection method, we observed that very few SNr units from control animals exhibit an oscillation in the 0.5–4 Hz range (2 of 85 units pooled across animals), whereas in each bilaterally dopamine depleted animal, 33–92% of units exhibited significant delta oscillations (117 of 226 units pooled) three days after depletion (Figure 3a). Without using the phase shift criterion, a much greater number of units were flagged as oscillating, particularly in control mice (28% of units vs 2% after phase shift correction, Figure 3b), despite these units having a nearly flat autocorrelation as in Figure 2c. To determine whether these oscillations remained stable at longer time points after depletion, we recorded from the SNr of unilaterally depleted mice, 2-4 weeks after depletion. We found that a significant proportion of SNr neurons still exhibited delta oscillations at these later time points (22–82% for each animal, 48 of 83 units pooled), suggesting that these oscillations are a stable feature of basal ganglia pathophysiology following dopamine depletion. A small number of units on the contralateral side of the lesion also exhibited delta oscillations (0–19% for each animal, 7 of 72 units pooled) (Supplemental Figure 1).

**Figure 3.**
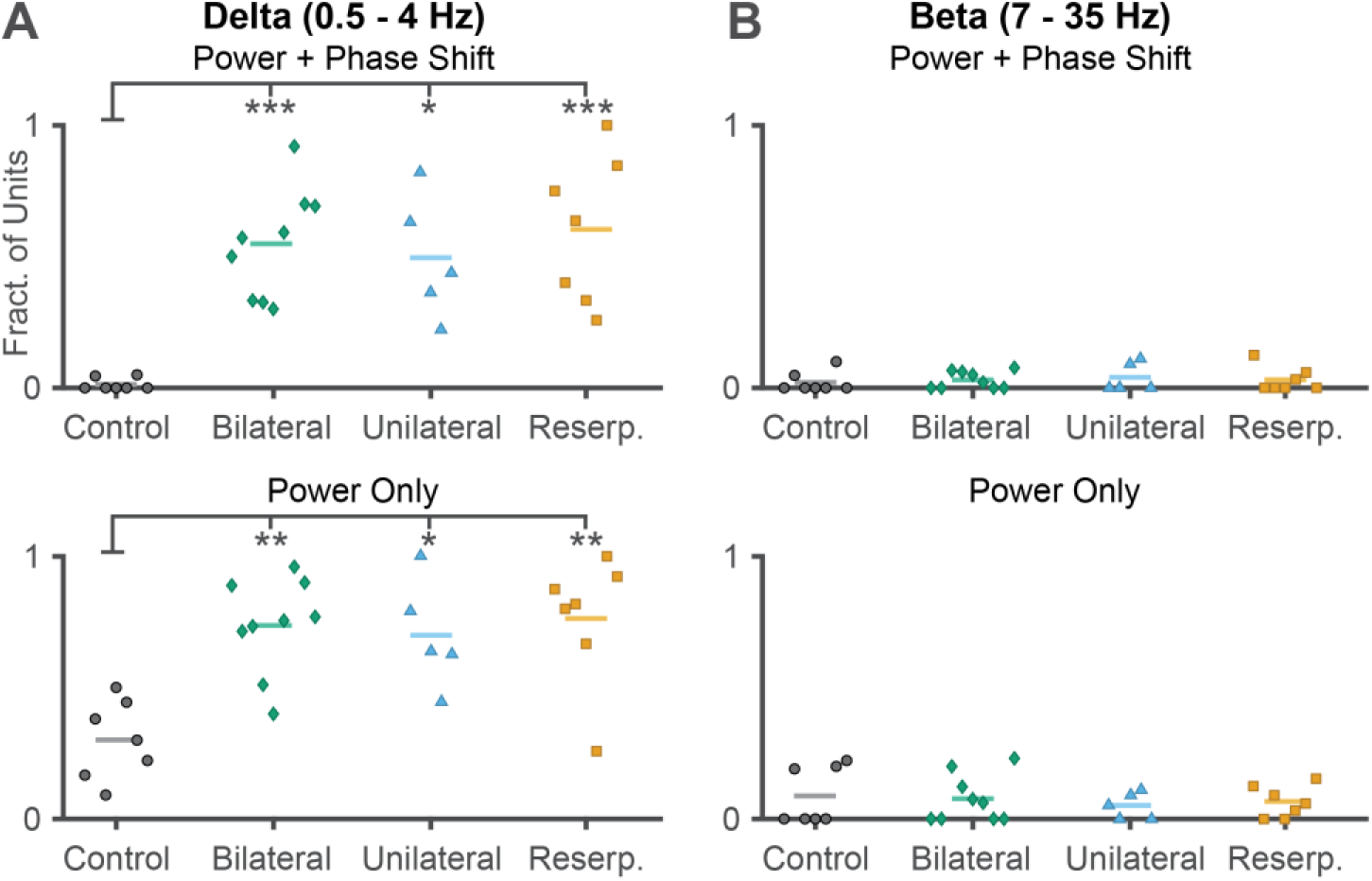
Dopamine depleted, but not control, SNr units exhibit phase-consistent delta oscillations, but no change in beta oscillations. Fraction of oscillating units from each animal in control conditions (black circles, n = 7) or various methods of dopamine depletion – bilateral 6OHDA (green diamond, n = 9), unilateral 6OHDA (blue triangle, n = 5), or systemic reserpine (orange square, n = 7). Lines indicate mean. **a.** Delta (0.5–4 Hz) oscillations detected using both PSD peak and low phase shift criteria. ANOVA: p = 5.206*10^-5^; bilateral: p = 9.506*10^-5^; unilateral: p = 0.00172; reserpine: p = 5.908*10^-5^, Dunnett’s post-hoc test. **b.** Same as **a**, but using only the spectral power criterion. ANOVA: p = 4.668*10^-4^; bilateral: p = 5.645*10^-4^; unilateral: p = 0.00601; reserpine: p = 5.794*10^-4^. **c**–**d.** Same as **a–b**, but for beta (7 – 35 Hz) oscillations. With phase shift, ANOVA: p = 0.8936; without phase shift, ANOVA: p = 0.8908

To ensure that delta oscillations were not merely an immune or inflammatory side effect of the injected toxin or cell death, we treated a cohort of animals intraperitoneally with reserpine, a compound that blocks the vesicular monoamine transporter 2 (VMAT2) complex from packaging monoamines into vesicles. This yielded a monoamine (including dopamine) depletion without any intracranial injection or cellular death and produced akinetic symptoms similar to those observed in bilateral 6-OHDA depleted mice. When we recorded three days after the start of daily reserpine injections, these animals exhibited a high proportion of slowly oscillating units in the SNr (33-100% for each animal, 74 of 119 units pooled), similar to bilaterally depleted animals (Figure 3a).

Given the prevalence of beta oscillations in the DD and PD literature, we sought to determine if these animals’ SNr units also exhibited beta oscillations. We defined a wide frequency range for beta oscillations, 7–35 Hz, which fully encompasses the definition of beta oscillations across humans and common model species (monkey and rat). We saw no increase in the fraction of beta oscillating units after any form of dopamine depletion, with or without our phase shift criterion (Figure 3c–d). Taken together, our results suggest that delta, not beta, oscillations are the primary oscillatory feature in SNr spike trains of awake, dopamine depleted mice.

### Oscillations predict DD severity and behavior better than other physiological measures of dysfunction

To understand how delta oscillations relate to the severity of dopamine depletion, we used an existing dataset of SNr recordings from mice gradually depleted to varying levels of dopamine loss through successive small injections of 6-OHDA (Willard et al., 2019). In this data, we looked at the relationship between an animal’s fraction of units exhibiting a delta oscillation and its level of dopamine neuron loss (as measured by striatal tyrosine-hydroxylase (TH) immunoreactivity). Performing a linear regression to predict %TH remaining from oscillation fraction showed a relatively strong (r^2^ = 0.5267) and significant (p < 0.01 from a bootstrapped 99% confidence interval, see Methods) relationship between dopamine loss and the fraction of oscillating units (Figure 4a).

**Figure 4.**
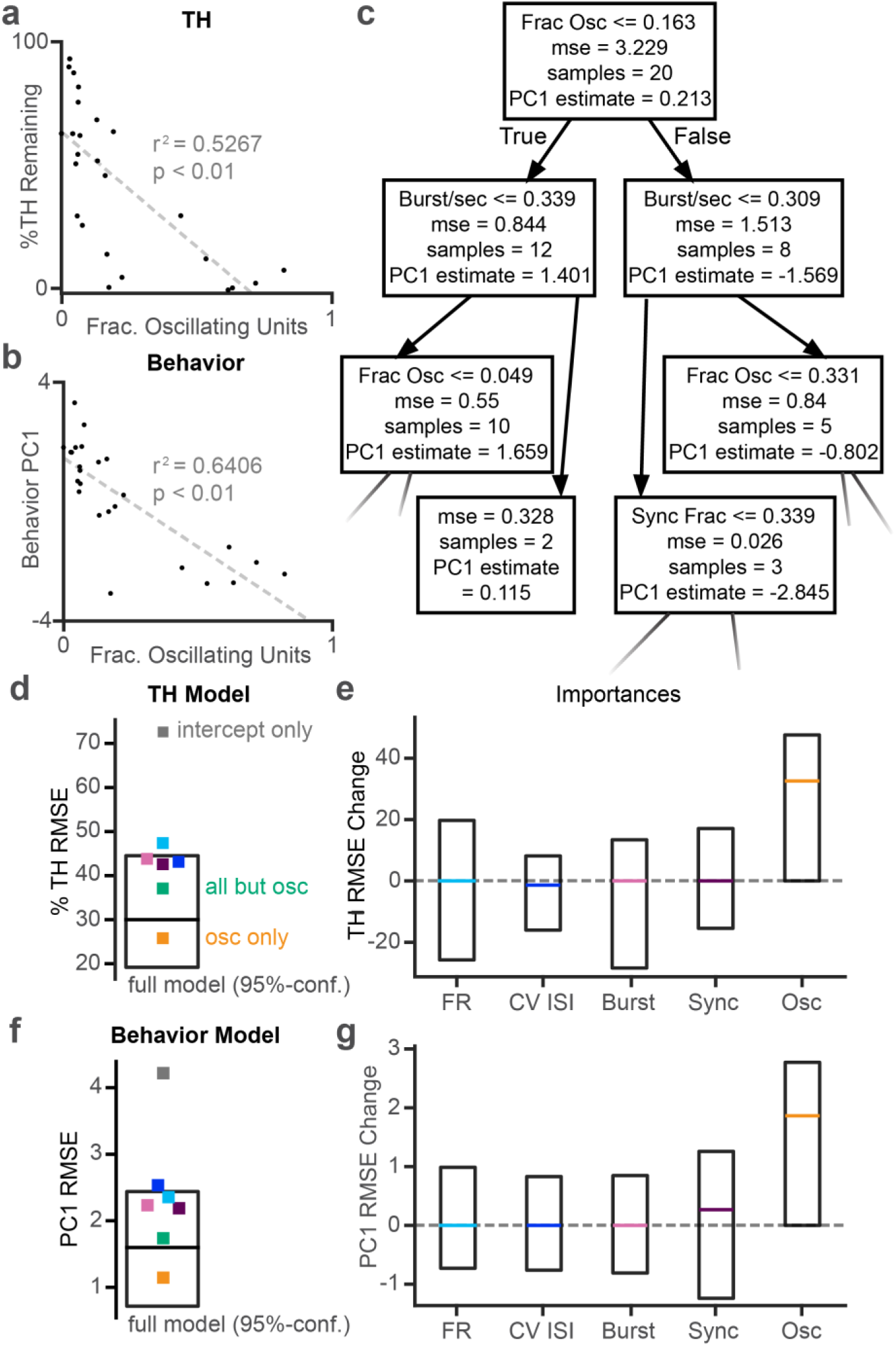
Oscillations predict severity of dopamine depletion. **a.** Scatterplot showing relationship between levels of remaining striatal TH and fraction of oscillating SNr units in animals (n=25) gradually dopamine depleted to different severities. Each dot denotes one animal, dashed line is the least squares fit. **b.** Same as **a** showing relationship between the first principal component (PC1) of several behavioral metrics (see methods, more negative indicates more dysfunctional) and the fraction of oscillating SNr units. **c.** The first three rows of one example decision tree predicting striatal TH from SNr neural properties (firing rate, irregularity, burstiness, synchronicity and fraction of delta oscillating units). **d.** A 95% confidence interval of MSE from 1,000 trees predicting TH. Each square is the MSE of the median model trained using a subset of parameters (grey: intercept-only, i.e. no parameters; light blue: firing rate; dark blue: CV of interspike intervals; pink: bursts/second; purple: mean synchrony across pairs; yellow: fraction of delta oscillating units; green: all parameters except fraction of delta oscillating units). **e.** Middle 95 percentile (box) and median (colored line, same color scheme as in **d**) of feature importances (permutation importance, see Methods) for each neural measure in the TH model computed from 1,000 trees. Dotted line indicates zero importance. **f–g.** Same as **d–e** for the model predicting PC1 of behavior.

Since striatal TH immunoreactivity is not a perfect indicator of parkinsonian symptoms, we also used these measures to predict motor behavior. Prior to *in vivo* recordings, these animals were given a series of behavioral tests to measure their mobility, dexterity, and strength (see Methods & Willard et al., 2019), and we performed principal component analysis on the results of these tests to get a single measure – the first principal component (PC1) – of their motor deficits. A linear regression predicting PC1 from the fraction of oscillating units illustrated a similarly strong and significant relationship (Figure 4b, r^2^ = 0.6406, p < 0.01).

Besides oscillations, many other neural measures in the basal ganglia have been suggested as correlates of DD severity – most commonly, changes in firing rate, firing regularity, burstiness, and synchrony between units. To see how delta oscillations compare to these measures in reliably predicting DD severity, we built a set of statistical models to predict %TH in each animal from five physiological parameters measured from single units in the SNr: 1) median firing rate, 2) median coefficient of variation (CV) of interspike intervals (ISIs), 3) median rate of bursts, as measured from the Poisson surprise test, 4) fraction of significantly synchronous pairs of units, and 5) fraction of units with significant 0.5–4 Hz oscillations. Due to the highly nonlinear relationship between the first four of these measures and DD severity (Willard et al., 2019), we performed a series of nonlinear regressions on this data by building 1000 decision trees from randomly selected sets of 20 (out of 25) animals, excluding the remaining 5 animals as a testing set for each tree (Figure 4c). We estimated a 95% confidence interval of mean squared errors (MSE’s) from these 1000 trees and showed that a tree built from these parameters predicts TH significantly better than a naive intercept-only model (Figure 4d).

To determine how each parameter informs the model, we shuffled the testing data for that parameter and calculated how much this loss of information increased the MSE of the model (the ‘importance” of that parameter). We then estimated 95% confidence intervals for the importance of each parameter (see Methods). The fraction of oscillating units was the only parameter whose confidence interval did not extend below zero (Figure 4e), suggesting that, when the model is built to include oscillations, they are the only parameter that provides reliably predictive information. In other words, while other parameters may provide information, that information is redundant when the fraction of oscillating units is known.

To confirm this in another manner, we rebuilt the models using the same cross-validated training and testing sets as above using only a single parameter at a time, or using all of the parameters except oscillations. The first four parameters fall outside (FR) or on the edge (CV ISI, Burst and Sync) of the full model confidence interval, but the model with all four parameters performs better than any individual parameter (Figure 4d), confirming results seen previously (Willard et al., 2019). However, the model built using only oscillations as a predictor is, on average, better than any other model including the combined parameter model, providing further evidence that other physiological parameters are not additionally informative when oscillations are considered.

Using the same procedure as above to predict PC1 of the animals’ behavior, we found very similar results to those predicting TH levels – namely, firing rate, irregularity, burstiness and synchrony provide some information in predicting behavior, particularly when considered together. However, when the fraction of oscillatory units is included in the model, it is the only important variable, and is significantly so, in predicting motor dysfunction (Figure 4f–g).

### Delta oscillations arise immediately following loss of MFB transmission or D2 receptor activation

The mechanism behind the observed delta oscillations is unclear, but they could arise due to a wide range of immediate biophysical changes in the basal ganglia after DD or emerge more slowly through plasticity or compensation. To determine this time course, we recorded from the SNr of healthy animals while acutely infusing lidocaine (a voltage-gated Na^+^ channel blocker) into the medial forebrain bundle (MFB), the same injection site for 6-OHDA in our other experiments, to quickly disrupt MFB transmission. We found that oscillations arose in the SNr within 2 minutes of the start of lidocaine infusion (before infusion ended) and waned within ten minutes after the end of infusion, mirroring the time course of akinesia observed during the experiment (Figure 5a-c). This result is consistent with the similarly rapid onset of slow oscillations produced by TTX infusion to the MFB under anesthesia (Galati et al., 2010) and demonstrates that low frequency oscillations arise in the SNr almost immediately after loss of MFB transmission, ruling out long-term mechanisms for their generation.

**Figure 5:**
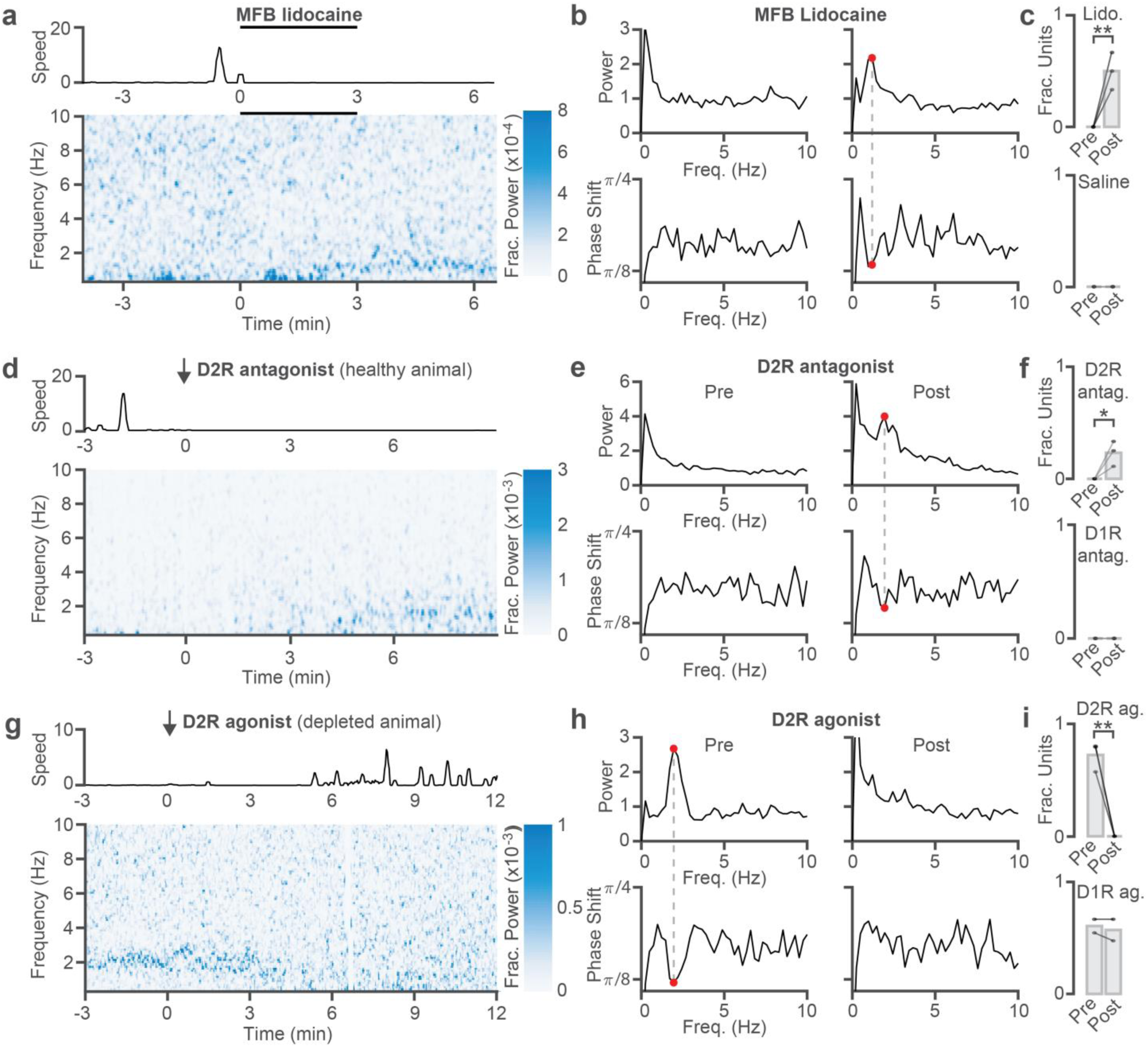
Acute manipulations of MFB signaling or D2-receptors modulate oscillations. **a.** Effects of lidocaine infusion into the MFB of healthy mice. Top: Speed of mouse on running wheel during lidocaine infusion (black bar). Bottom: Spike spectrogram of an example SNr unit during the same infusion as above. **b.** Top: PSDs from the same unit before (left) and after (right) lidocaine infusion. Bottom: Phase shift plots corresponding to the above PSDs. A dashed line from the detected oscillation in the right PSD (red dot) connects to the same frequency in the corresponding phase shift plot **c.** Fraction of oscillating units from all animals before and after lidocaine (top, n = 3, p = 0.00219) or saline (bottom, n = 2, p = 1.000) infusion into the MFB. Each dot is one animal, bars indicate mean, and lines connect the same animal before and after infusion. **d**–**f.** Same as **a–c**, but for systemic injection of a D2R antagonist (raclopride, n = 3, p = 0.0233) compared to a D1R antagonist (SCH233890, n = 2, p = 1.000) **g**–**i**. Same as **d**–**f**, but for systemic injection of a D2R agonist (quinpirole, n = 3, p =8.686*10^-4^) compared to a D1R agonist (SKF81297, n = 2, p = 0.7455) in dopamine depleted animals.

To determine whether the loss of dopamine signaling is causal to the onset of delta oscillations, we recorded from the SNr of healthy animals before and during the systemic injection of a D1-receptor (D1R) antagonist (SCH233890) or a D2-receptor (D2R) antagonist (raclopride). While both drugs caused reduced movement on the wheel, only the D2R antagonist led to the development of oscillations in the SNr (Figure 5d–f). We then performed the converse experiment, injecting a D1R agonist (SKF81297) or D2R agonist (quinpirole) systemically into bilateral DD animals. Similarly, while both led to highly increased motor activity (though highly dyskinetic in the case of D1 agonism), only the D2R agonist injection attenuated delta oscillations in the SNr (Figure 5g–i). This suggests that low frequency oscillations are mediated purely due to a loss of action on D2Rs and are not affected by D1Rs.

### Delta oscillations are a feature of dopamine depletion throughout the indirect pathway

Since the indirect pathway of the basal ganglia is a primary location of D2R-expressing neurons, we posited that oscillations may also be present elsewhere in the indirect pathway. We thus recorded from healthy and dopamine depleted globus pallidus externa (GPe) (Figure 6a) and subthalamic nucleus (STN) (Figure 6c), two reciprocally connected nuclei in the indirect pathway that both project heavily to SNr. We found a similar pattern of oscillatory activity across units in the GPe (40–80% of units in each animal, Figure 6b) and STN (15–70% of units in each animal, Figure 6d) after dopamine depletion, whereas only 1 of 111 total GPe and 1 of 63 STN units exhibited oscillations in the healthy state.

**Figure 6:**
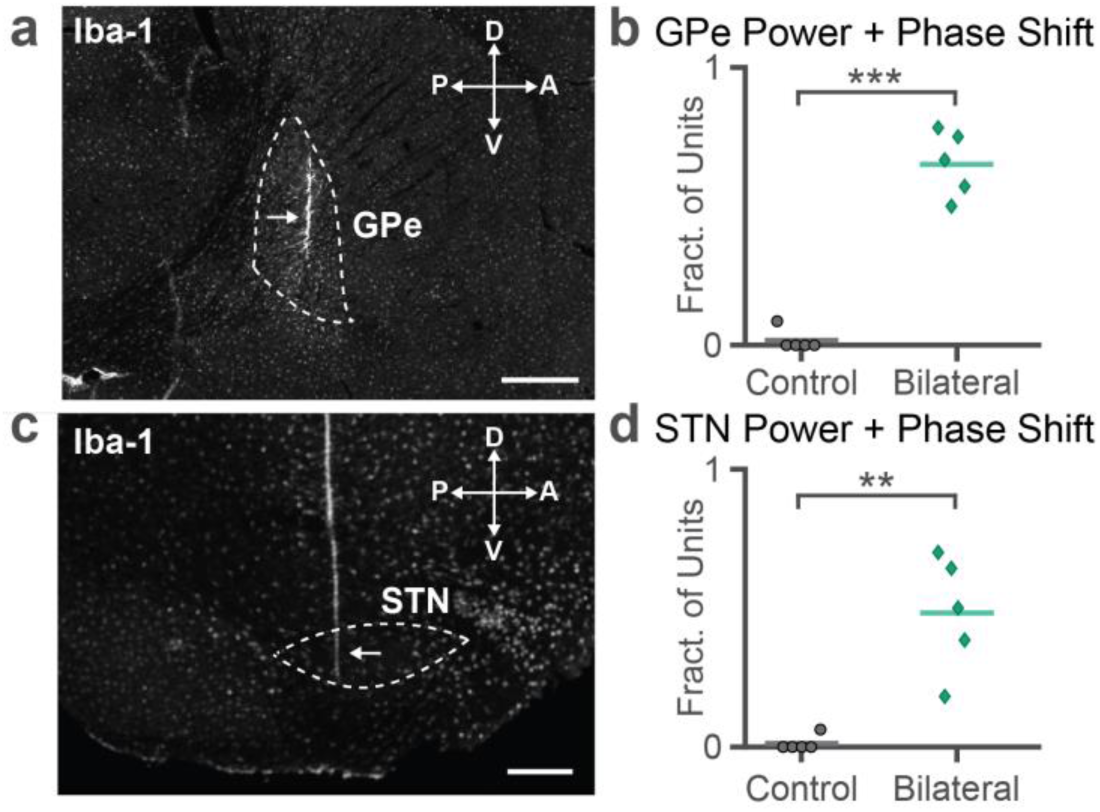
Delta oscillations pervade the dopamine depleted, but not healthy, indirect pathway. **a.** IBA immunofluorescence showing example probe locations in GPe. Dotted line indicates approximate location of GPe, arrow indicates probe location. Scale bar = 500 µm. **b.** Fraction of oscillating units from each animal in control (black circles, n = 5), or bilateral 6OHDA (green diamond, n = 5) animals in GPe (p = 3.847*10^-6^, two-sample t-test). **c**–**d.** Same as **a–b** targeting STN (p = 0.00106, both control and bilateral n = 5)

### Two populations of delta oscillating units in SNr both lead oscillations in motor cortex

Previous literature suggests that oscillations in the dopamine depleted basal ganglia arise due to input from oscillating neurons in motor cortex (M1) under anesthesia (Tseng, Kasanetz, Kargieman, Riquelme, et al., 2001). However, since we have shown that these oscillations arise from antagonism on D2R’s, a receptor more prevalent in the basal ganglia than M1, a possible alternative in awake animals is that these oscillations arise first in the basal ganglia and then entrain M1.

To distinguish between these possibilities, we sought to characterize oscillations in the M1 of DD animals and determine the phase lag between M1 and SNr oscillations. We recorded an electrocorticogram (ECoG) in M1 while simultaneously recording from single units in SNr (Figure 7a). Compared to healthy controls, the M1 ECoG of DD animals exhibited a large increase in delta oscillations and reduction in theta (4–7 Hz) oscillations, which are typically seen in the cortex of healthy mice (Tort et al., 2018) (Figure 7b–c).

**Figure 7:**
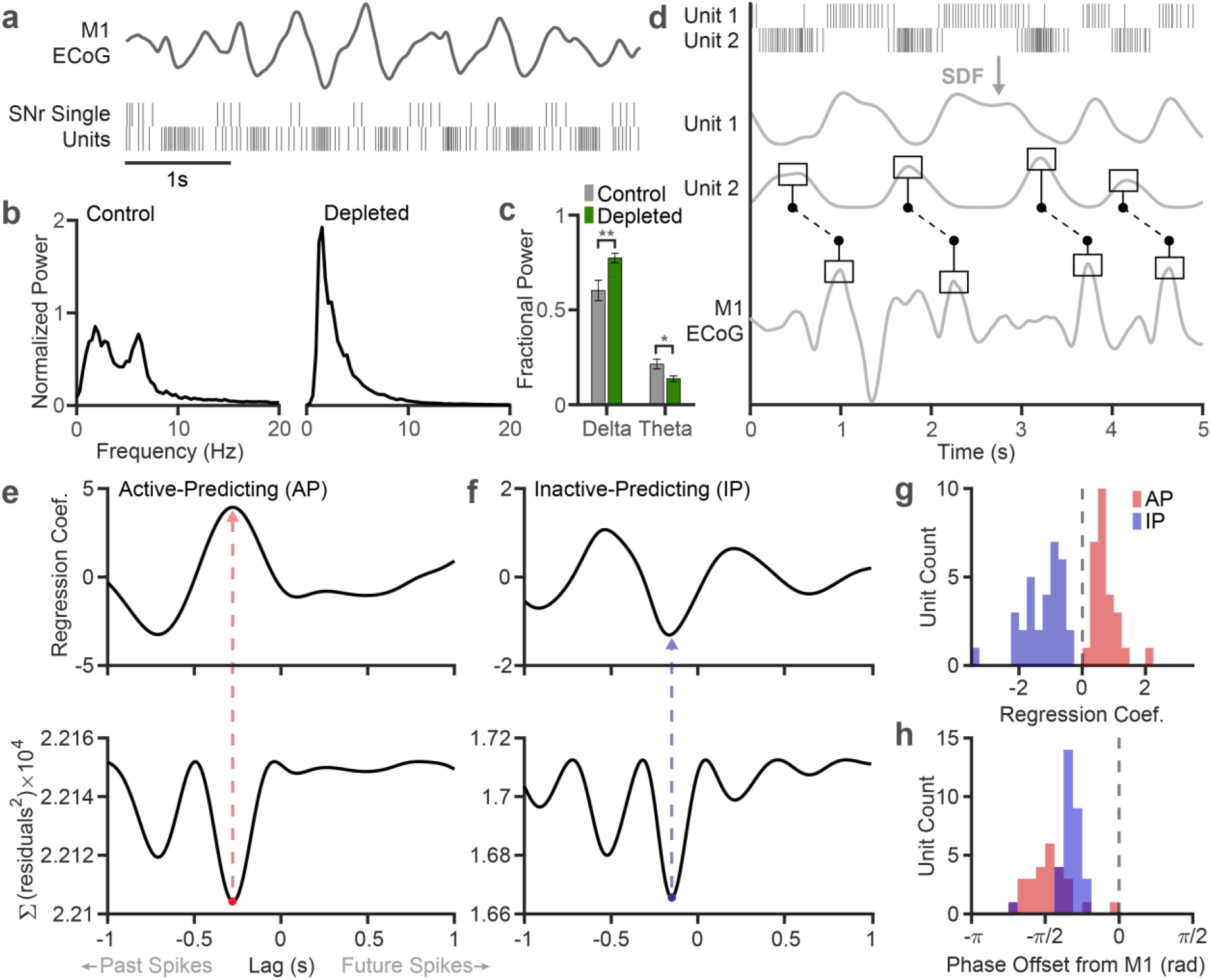
Delta oscillations define two SNr populations which both lead oscillations in M1. **a.** Example simultaneous M1 ECoG and spike trains from two SNr units exhibiting coherent oscillations. **b.** Example M1 ECoG power spectra from control (left) and bilaterally depleted (right) animals. Power spectra were normalized to their total 0.5-100 Hz power and multiplied by 1000 for visualization. **c.** Fractional delta and theta band power in M1 ECoG across all control (n = 8) and acutely depleted (n = 9) animals. Bars indicate mean, error bars indicate standard error (p = 0.00818 for delta, 0.0173 for theta, two-sample t-test test). **d.** Example data demonstrating SNr predicting M1. Top: 5 second rasters from two simultaneously recorded SNr units. Middle: spike density functions (SDF) of the above SNr rasters. Bottom: Simultaneously recorded M1 ECoG. Lines between the bottom two panels illustrate M1 exhibiting peaks at a consistent time lag after the peak of an SNr SDF, even amidst variance in oscillation period length. **e.** Example regression results predicting M1 ECoG from an “active-predicting” (AP) SNr unit. Top: Regression coefficients for each individual lag. Negative lag corresponds to SNr oscillations leading M1. Bottom: MSE of regression results using each lag. The red dot indicates that the model using that lag significantly outperforms an autoregressive model of the ECoG (F-test, p < 0.05 correcting for multiple lag comparisons). The dotted line to the upper panel lands at a peak in the coefficients, defining the unit as “active-predicting”. **f.** Same as **e** for an “inactive-predicting” (IP) SNr unit, whose significant lag is labeled in blue. **g.** Summary histogram of regression coefficients from all oscillating SNr units recorded simultaneously with M1 ECoG (n = 59). Counts are colored as in **e–f** based on their regression coefficients (red: positive, blue: negative, dashed line at zero), which define their type (AP or IP). **h.** Same units colored as above grouped by the phase offset at which they best predict the M1 ECoG (as in **e–f**, negative phase offsets correspond to SNr oscillations leading changes in M1).

Determining the relationship between two oscillating signals from their phases is a difficult task – if the phase of one perfect oscillator slightly leads that of a second perfect oscillator, it is impossible to distinguish whether the first leads the second at a short lag or if the second leads the first at a long lag. However, neural oscillations do not match the activity patterns of perfect oscillators, but in fact have profiles that vary across periods and highly varying period lengths that are merely centered on a range of values. We can leverage this fact to make predictions about the relative timing of SNr and M1 (Figure 7d).

To quantify this relation, we performed a series of Granger causality regressions, which make no assumptions on the periodicity of the signals. Rather, they simply attempt to predict changes in M1 ECoG based on its own history (the null, autoregressive model) or by additionally including SNr spiking information from a single unit. For each unit, we computed 201 separate models predicting M1, each using SNr spiking information at a different lag between −1 (i.e., past spikes) and +1 seconds (i.e., future spikes). Aligning the lag coefficients of the models for a single unit illustrates a periodicity in their values that matches the oscillation period (Figure 7e-f).

We computed the mean squared error (MSE) of each model at each lag and considered the lag that minimized MSE. To quantify whether this model significantly outperforms the purely autoregressive ECoG model, we performed an F test on the two models, correcting for multiple lag comparisons (Figure 7e-f). We find that 51 of 63 of oscillating units in SNr predicted changes in the ECoG significantly better than the null autoregressive model, suggesting that there is significant correlation between SNr and M1 at a consistent time lag.

When analyzing the regression coefficients at these significant lags, we found a clear bimodal distribution of units determined by whether the active or inactive phase of their spike oscillation predicted positive deflections in M1. We term these types “active-predicting” (AP) units, which make up approximately 47.1% of ECoG-locked units (38.1% of oscillating units, 34.3% of all analyzed units) and “inactive-predicting” (IP) units (Figure 7g), which make up the remaining 52.9% of ECoG-locked units (42.9% of oscillating units, 38.6% of all analyzed units). We see further evidence of these two distinct populations through cross correlation analysis of SNr unit pairs (Supplementary Figure 2).

When clustering units based on their phase lag relative to M1, SNr units also organize into a bimodal distribution, with one mode dominated by AP units and the other by IP units. (Figure 7h). Critically, all significant lags were negative – that is, SNr spikes from both populations of SNr units consistently predicted future changes in the ECoG, but not the inverse (Figure 7h). The relative timings of these signals suggest an order in which oscillations propagate through the SNr and cortex - AP neurons enter their active phase (increase firing), then IP neurons enter their inactive phase (decrease firing or pause), and finally M1 enters its active phase. These results suggest a consistent timeline of oscillatory dynamics by which two oscillating populations in SNr both dynamically predict M1 activity.

### M1 is not required for delta oscillations in SNr

The results of our regression analysis suggest that oscillations in SNr are not caused by M1, but rather that oscillations in the SNr precede and predict those in M1. To test this hypothesis, we performed M1 aspiration lesions in dopamine depleted mice (Figure 8a) and recorded from the SNr. SNr units in the DD + M1-lesioned mice had similar oscillations to those DD mice without M1 lesions (Figure 8b). These mice had a significantly higher fraction of oscillating units than control animals, but there was no difference between dopamine depleted animals with or without an M1 lesion (Figure 8c). These results provide additional evidence that M1 is a recipient, not the source, of delta oscillations in DD.

**Figure 8:**
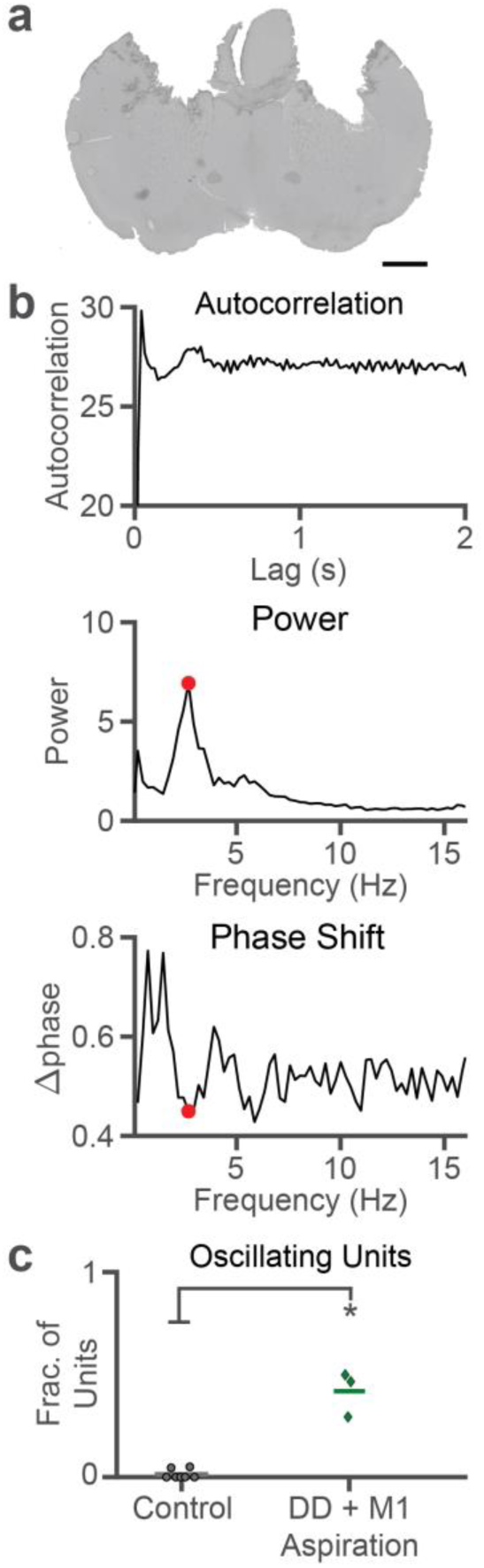
M1 lesion does not disrupt oscillations in SNr. **a.** Example coronal slice from an M1 lesioned animal. Scale bar = 1 mm. **b.** Autocorrelation (top), PSD (middle) and phase shift (bottom) for an example SNr unit exhibiting a delta oscillation in an M1-lesioned, dopamine depleted animal. **c.** Fraction of oscillating units in SNr for each animal in control (black circle, n = 7) and bilaterally dopamine depleted with M1 lesion (dark green diamond, n = 3) conditions (p = 1.1478*10^-6^, two-sample t-test.)

## Discussion

In this paper, we have demonstrated that delta (0.5–4 Hz), not beta (7–35 Hz), oscillations are the predominant oscillatory feature in basal ganglia neurons in awake, dopamine depleted mice, and that the fraction of units exhibiting these oscillations is a good marker of dopamine loss and motor deficits. These results are consistent with data from the human PD literature demonstrating that delta oscillations are the dominant or only oscillatory feature in some PD patients (Du et al., 2018; Levy et al., 2002). We further show that these oscillations arise from a loss of action on D2 receptors and that, contrary to conclusions drawn from anesthetized experiments, motor cortex is not required for their generation but rather follows the oscillations evident in the basal ganglia.

### A novel method to distinguish oscillations from noise

Although several studies demonstrate the presence of delta oscillations in the LFP (Levy et al., 2002; Priori et al., 2004) and single units (Steigerwald et al., 2008; Du et al., 2018; Zhuang et al., 2019) of PD patients, many more studies ignore oscillations in this band completely. Difficulties in detecting these oscillations may contribute to this lack of attention. Most studies examining oscillations in PD patients investigate the LFP, not individual spiking units, and the intrinsic low frequency noise of LFP signals makes reliably detecting oscillations in the delta range difficult. Even when it is possible to record from single units, we have demonstrated that low frequency noise can disrupt these spiking signals as well.

To reliably detect low frequency spike oscillations in awake animals, we have introduced phase shift as a novel detection technique which utilizes phase information typically discarded from the Fourier transform. Phase shift measures the local stationarity of a signal composed primarily of one frequency – a perfect sine wave would have zero phase shift and high power, but a sine wave with a phase that randomly advances would have high phase shift while maintaining high power. This measure can distinguish our signal of interest – a single oscillatory signal that shifts in phase only gradually or rarely – from low frequency pink noise, a phenomenon that is not restricted to a single frequency, for which phase components measured at individual frequencies may shift rapidly between adjacent windows.

### Relationship to previous studies on PD oscillations

In PD research, much of the oscillation literature has focused on the beta band (Hammond et al., 2007; Jenkinson & Brown, 2011). Here, we demonstrate dopamine loss and PD-like symptoms in mice without the presence of beta oscillations, and we have previously demonstrated their absence in LFP signals in awake mice as well (Willard et al., 2019). Indeed, to our knowledge, no study has demonstrated the presence of beta oscillations in mouse models of PD. Instead, this study suggests that delta oscillations are an important signal in the dopamine depleted basal ganglia and may cause parkinsonian dysfunction instead of or (in patients or other animal models) alongside beta oscillations.

The low frequency oscillations that we observe resemble those seen in anesthetized mice and rats, although oscillations in awake settings are generally noisier. By performing these experiments in awake mice, this study rules out concerns that oscillations in the basal ganglia are simply entrained by anesthesia-induced oscillations from cortex (Tseng, Kasanetz, Kargieman, Riquelme, et al., 2001; Belluscio et al., 2003) or by artificial respiration devices (Ruskin et al., 2002). Instead, we see that oscillations in the basal ganglia arise even during wakefulness and in fact lead and predict oscillations in M1. While we can rule out one causal direction (M1 -> SNr), it is difficult to know whether SNr entrains M1 directly or if both SNr and M1 are entrained by a common source.

By referencing SNr oscillations to M1, we distinguish two populations of oscillating SNr neurons. These populations and how they are defined mimic the Type-A (TA) and Type-I (TI) populations observed in GPe whose discharge is high and low, respectively, during the active phase of M1 oscillations (Mallet et al., 2008). Active-predicting (AP) and inactive-predicting (IP) SNr neurons are a very close analog to TA and TI GPe neurons, respectively, except for two differences. First, the granularity of our regression analysis illustrates that AP and IP neurons are not simply active or inactive during the active phase of the M1 oscillation, but begin discharging (AP) or pausing (IP) 150–250 ms before the active component of the M1 oscillation. To our knowledge, a precise timing analysis of TA and TI neurons with M1 oscillations has not been performed to determine if a similar phenomenon occurs in GPe. Second, SNr AP and IP neurons are approximately equal in number, whereas TI neurons are the prevailing population in GPe (72% TI, 17% TA) (Mallet et al., 2008). These populations of GPe neurons were later shown to have anatomical (Corbit et al., 2016; Mallet et al., 2012), genetic (Abdi et al., 2015), and functional (Gage et al., 2010; Mallet et al., 2016) differences, forming the prototypic (TI) and arkypallidal (TA) populations. AP and IP neurons may exhibit such differences as well, although these analyses would be beyond the scope of our study.

### Mechanisms of generation: insight from D2 receptors

A previous study demonstrated that delta oscillations in anesthetized mice arise immediately after loss of dopamine signaling through the MFB (Galati et al., 2010), a finding that we have replicated here in awake mice. This fast onset (<2 minutes) contrasts with the typical longer timescale associated with beta oscillations in DD (Mallet et al., 2008). We further show that these oscillations arise due to a loss of D2R activation and can be ablated in already dopamine depleted animals through D2R agonism. It is unclear where the D2 receptors responsible for this ablation are located, but the high density of D2R’s in the striatum make it a strong candidate. Lack of D2R activation causes a wide array of biomolecular changes within D2R-expressing neurons, including the opening of NMDA (Higley & Sabatini, 2010; Wang et al., 2012) and L-type calcium channels (Hernández-López et al., 2000), which have been shown to be involved in membrane potential and calcium oscillations, respectively, in other circuits (Guertin & Hounsgaard, 1998).

In addition to striatum, another candidate for the generation of delta oscillations in DD is the STN-GPe loop. While often associated with beta oscillations (Mallet et al., 2008; Nevado-Holgado et al., 2014; Pavlides et al., 2012; Wei et al., 2015), this loop was originally implicated in generating much lower frequency oscillations (0.8 – 1.8 Hz) in cultured neurons (Plenz & Kital, 1999), a phenomenon that has been demonstrated subsequently in computational models (Terman et al., 2002; Modolo et al., 2008). The slow rates associated with the dynamics of T-type calcium channels and of some after-hyperpolarization currents have been shown to contribute the generation of oscillations and could explain the low frequency of these oscillations as well (Devergnas et al., 2015).

### Relationship between oscillations and motor dysfunction

Of those studies that examine low frequency oscillations in PD patients, many consider only their relationship to tremor, seeing both positive and zero correlation with EMG signals during tremor bouts (Hurtado et al., 1999; Du et al., 2018). No study, to our knowledge, has investigated low frequency oscillations in relationship to other PD symptoms. Here, we have established a strong relationship between delta oscillations, dopamine loss, and akinetic dysfunction in mice. Further research and re-examination of existing patient data could elucidate a role for delta oscillations in predicting or causing PD motor deficits in humans.

While we cannot demonstrate a causal link between oscillations and motor dysfunction in this study, it is notable that the emergence of delta oscillations in the SNr from multiple experimental manipulations is consistently paired with a time-locked and commensurate reduction in motor activity. These results suggest a reappraisal of delta oscillations as a potential cause or marker of motor dysfunction in Parkinson’s disease patients that could be an underappreciated target for PD therapies.

## Methods

### Animals

All experiments were conducted in accordance with guidelines from the National Institutes of Health and with approval from the Carnegie Mellon University Institutional Animal Care and Use Committee. Male and female mice on a C57BL/6J background aged 8-15 weeks were randomly allocated into experimental groups (e.g. Control, Bilateral 6OHDA, Reserpine, etc.), except insofar as to ensure that male and female mice were both represented in every group.

### Stereotaxic surgery

#### Headbar implantation

Animals were anesthetized with 20 mg/kg ketamine and 6mg/kg xylazine and placed in a stereotaxic frame (Kopf Instruments). Anesthesia was maintained throughout surgery with 1.0-1.5% isoflurane. All coordinates were measured in mm with AP and ML measured from bregma and DV relative to the dural surface. The scalp was opened and bilateral craniotomies (for later probe insertion) approximately 1.5 x 1.5 mm in size were drilled over SNr (AP: −3.00, ML: ±1.50), GPe (AP 0.00, ML: ±2.12), or STN (AP: −1.70, ML: ±1.52). A custom-made copper or stainless steel headbar was affixed to the mouse’s skull with dental cement (Lang Dental). A well of dental cement was then built around the exposed skull (see *in vivo recordings*) and filled with a silicon elastomer.

#### Dopamine depletion

A hole was drilled on one (for unilateral) or both (for bilateral) hemispheres of the skull over the medial forebrain bundle (MFB, AP: −0.80, ML: ±1.10). A unilateral infusion cannula (PlasticsOne) was slowly lowered into the brain 5mm below the dura. 1 µL of 5 µg/µL 6OHDA (Sigma-Aldrich) or 0.9% saline was injected over the course of 5 minutes with a GenieTouch Hamilton syringe pump (Kent Scientific). The infusion cannula was left in place for 5 minutes post-injection before being slowly retracted. For animals undergoing bilateral depletion, this process was repeated on the opposite hemisphere.

#### Cannula implantation

For experiments involving acute drug infusion into the MFB or gradual dopamine depletion with 6OHDA, a bilateral guide cannula (Plastics One) was implanted (same coordinates as dopamine depletion) using dental cement (Lang Dental) and a dummy cannula was placed in the guide. Before infusion, the dummy was replaced with an infusion cannula and attached to the same Hamilton syringe pump as above. Gradually depleted animals were infused with 1 µL of 0.75 µg/µL 6OHDA every 5 days (See Willard et al 2019 for full details).

#### ECoG connector implantation

For experiments involving electrocorticogram (ECoG) recordings, a male gold connector (Ampityco Electronics) was soldered to a stainless steel wire, and the connector was gently lowered above left or right motor cortex (M1, AP: +1.40, ML: ±1.00) such that the wire touched the dural surface then secured in place with dental cement (Lang Dental).

#### Aspiration lesions

For experiments involving M1 lesion, a craniotomy was drilled bilaterally over M1 (AP 0.0-2.5, ML 1.0:2.5) and the dura was removed. Using a 20-gaugse suction tube (Miltex) attached to a vacuum source, we aspirated cortex to a depth of 2.5 mm across the craniotomy and under portions of the remaining skull, periodically lightly rinsing the area with saline. We filled the lesioned space with triple antibiotic (bacitracin, neomycin, polymyxin) before sealing the craniotomy with a silicon elastomer (Smooth-On)

#### Post-operative care

Upon completion of surgery, animals were injected subcutaneously with 0.5 mg/kg ketofen and placed inside their cage half on/half off a heating pad to recover. Dopamine depleted animals were supplied with trail mix and moistened food to maintain weight and hydration, in addition to their usual food pellets and water bottles, and animals were tracked regularly to ensure proper health and weight.

### Drugs

In addition to the drugs used above during surgery, animals were given the following drugs (Sigma-Aldrich, except when specified) dissolved in 0.9% saline (except when specified). For reserpine depletions, animals were injected i.p. daily for three days with 5 mg/kg reserpine in 2% acetic acid (diluted in 0.9% saline). For recordings involving dopamine agonists and antagonists, animals were injected i.p. during recording with either 0.4 mg/kg SCH22390, 3 mg/kg raclopride. 1 mg/kg SKF81297 (Tocris Biosciences), or 3 mg/kg quinpirole. Acute infusions into the MFB used 2% lidocaine.

### In vivo recordings

Mice were head-fixed atop a free-running wheel (Heiney et al., 2014). After acclimation to head-fixation for ten minutes, the silicon elastomer was removed and craniotomies were cleaned with saline. Using a micromanipulator (Sutter Instruments), a linear microelectrode probe with sixteen channels spaced 50 µm apart (NeuroNexus) was lowered into the craniotomy at the coordinates listed above for SNr, GPe or STN. After the initial lowering, a ground wire was placed in saline in the dental cement well on the skull. Once the top of the nucleus (SNr, −4.0mm, GPe: −3.60mm, STN: −4.00mm from the top of the brain) was found and high firing rate units were observed, the probe was held stable for at least ten minutes prior to recording. Spiking (bandpass filtered for 150-8000 Hz, sampled at 40 kHz) and local field potential (bandpass filtered to 0.5-300 Hz, sampled at 1 kHz) recordings were collected through an OmniPlex amplifier (Plexon, Inc.) with common median virtual referencing. After recording for at least three minutes, the probe was lowered to explore the full dorsal-ventral extent of the nucleus. Simultaneous to these recordings, the mouse’s walking speed on the wheel was recorded using an optical mouse and fed to a TTL-pulser which was connected to the OmniPlex amplifier analog input. For ECoG recordings, the gold implant was connected to a headstage with a ground wire in saline on top of the skull. The headstage was connected to an amplifier (A-M Systems) with 1000x gain and 0.1–500 Hz bandpass filtering and this amplifier was connected to the OmniPlex amplifier analog input.

### Histology

After recording, animals were sacrificed and perfused with 4% paraformaldehyde (PFA). The brain was extracted from the skull and stored in PFA for 24 hours then moved to a 30% sucrose solution for at least 24 additional hours. Tissue was sectioned using a freezing microtome (Microm HM 430; Thermo Scientific) and primary antibody incubations were performed on these sections at room temperature for 24 hours. A tyrosine-hydroxylase (TH) antibody (rabbit anti-TH, 1:1000; Pel-Freez) was used to confirm successful dopamine depletion in 6OHDA-depleted animals; animals required at most 15% TH fluorescence compared to controls on both hemispheres (for bilateral 6OHDA injection) or the contralateral hemisphere (for unilateral 6OHDA injection) to be considered for analysis. and an Iba1 antibody (rabbit anti-Iba1) for microglia activation was used to confirm probe location and guide cannula placement in animals undergoing infusion during recording. Epifluorescent images were taken at 10x magnification (Keyence BZ-X) and outlines of nuclei of interest were overlaid on the images (from Paxinos Mouse Brain Atlas in Stereotaxic Coordinates, Second Edition).

### Data pre-processing

Spikes were manually sorted into single units using Offline Sorter (Plexon). For classification as a single unit, the following criteria were set: 1) principal component analysis of waveforms generated a cluster of spikes significantly distinct from other unit or noise clusters (p < .05), 2) the J3-statistic was greater than 1, 3) the Davies-Bouldin statistic was less than 0.5, and 4) fewer than 0.15% of ISI’s were less than 2ms. In the case where a unit was lost during recording, it was only used in analysis for the time period when its spike cluster satisfied these criteria, and only if its cluster was present for at least three minutes. Data were then imported into MATLAB (MathWorks) in which all further analysis was performed using custom code except when specified.

Since units must fire quickly enough to exhibit an oscillation, only units with a firing rate greater than 5 Hz (over 95% of sorted units) were considered for analysis. As ECoG signals were occasionally corrupted for short time windows, generally due to muscle activity, we visually determined a noise threshold for each recording and zeroed any length of signal within 250 milliseconds of any data point whose absolute value exceeded that threshold. ECoG signals were then delta (0.5–4 Hz) bandpassed using a 2^nd^ order Butterworth filter.

### Oscillation detection and visualization

#### Renewal-Corrected Power Spectrum

For each unit, we downsampled its spike train to 1 kHz and split it into segments of 2^12^ ms, advancing from one segment to the next with time step size *Δs* = 2^9^ ms. For each segment, we calculated its interspike interval (ISI) probability distribution, *P*_0_(*t*). We calculated 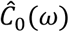, the theoretical power spectral density (PSD) of a renewal process defined by *P*_0_(*t*) scaled by the number of spikes in the segment:

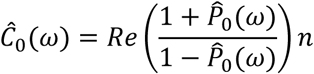

where *Re*(*x*) indicates the real part of x, 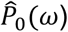 indicates the Fourier transform of the ISI distribution in appropriate frequency units, and n is the number of spikes in the segment. This is a variant of a method presented previously for calculating 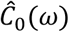 analytically rather than approximating it through Monte Carlo shuffling simulations (Rivlin-Etzion et al., 2006).

We next calculated an estimate of the PSD of the spike train in that segment:^1^

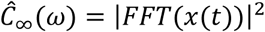

where x(t) is the mean-subtracted spike train in the segment, FFT is the fast Fourier transform (MATLAB function *fft*) and vertical bars indicate absolute value. Finally, we normalized this estimate to achieve the renewal-corrected PSD of a single segment:

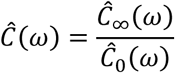

and averaged 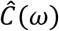 values across segments to obtain the renewal-corrected PSD. All PSD’s in this study have undergone this renewal-correction, but are simply referred to as PSD’s for brevity.

#### Phase Shift

For the kth time segment, we calculated the uncorrected phase 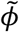 at each frequency:

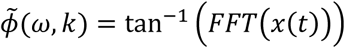

and made the following correction such that the phase of each frequency is defined relative to the start of the recording rather than the start of the segment:

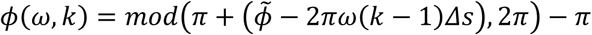

where mod is the modulus operator and *Δs* is the time step between adjacent segments (here, 2^9^/1000 seconds). In other words, for each frequency, imagine a perfect oscillator with zero phase at the start of the recording. For each segment, we determined what phase this oscillator would reach at the start of the segment and defined that phase to be zero for that segment. This correction ensures that a perfect oscillator would have the same corrected phase *ϕ* for every segment.

After computing the corrected phase of all segments, we approximated the time derivative *ϕ*_*s*_(*ω, k*) by computing the difference of phase across successive time steps and averaged over each difference to obtain the average absolute rate of phase shift:

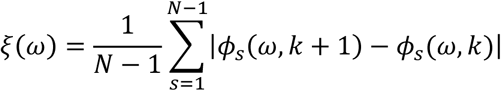

where *N* is the number of segments. For brevity, we refer to *ξ*(*ω*) as the phase shift.

#### Oscillation Detection

We detected oscillations in a two-step process by first seeking frequencies with high power and then determining whether these frequencies also had low phase shift.

To determine whether a unit reached statistically significantly high power at a particular frequency, we found each local maximum of 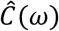, defined as a value higher than its three neighbors on both sides, within the band 0.5–4 Hz (or 7–35 Hz for detecting beta oscillations). We then estimated a 99% confidence interval of renewal-corrected power from the region of 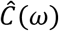 between 100 and 500 Hz, correcting for multiple comparisons (Bonferroni correction) of all frequencies in the band of interest. A peak of 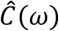 was considered significant if it fell above this confidence interval.

As our second step, we determined if any frequency detected in the previous step had a significantly low phase shift. We estimated a 95% confidence interval of phase shifts from the region of *ξ*(*ω*) between 100 and 500 Hz, correcting for multiple comparisons (Bonferroni correction) if multiple frequencies were detected from the PSD. We concluded that an oscillation was present at a frequency with significant power if the phase shift at that frequency fell below this confidence interval.

#### Spike Spectrograms

For time frequency analyses, the process outlined under *Renewal-Corrected Power Spectrum* was modified to use segments of length 2^13^ ms with 2^11^ ms overlap to improve visualization. Rather than averaging over segments, the resultant matrix was smoothed with a 3×3 2-D gaussian filter and plotted as a normalized heatmap (MATLAB function *imagesc*). Due to the loss of fine frequency resolution at low frequencies, this procedure was only used on spike trains in which an oscillation was detected in the previous procedure.

### Neural Measures

Beyond oscillations, we investigated several other neural measures – firing rate, firing variability, bursts and synchrony. A unit’s firing rate was defined as its number of spikes divided by the total time of recording. Variability was measured as the coefficient of variation (standard deviation divided by mean) of a unit’s interspike intervals. Bursts were quantified using the Poisson surprise algorithm (Legendy & Salcman, 1985) with a surprise threshold of 5, initial firing rate threshold of 200% of baseline calculated over the entire recording, and removal of any burst with fewer than 3 spikes.

To determine if two units were synchronous, we used the method and parameters outlined in Willard et al. 2019, which determines the fraction of synchronous spikes above chance after correcting for nonstationarity in a unit’s firing rate (Willard et al., 2019). In brief, we windowed both spike trains into 12-second segments and zeroed the first and last four seconds of the segment taken from the second spike train. We performed cross-correlation with a maximum lag of four seconds. Since this maximum lag is equal to the length of time zeroed on the second spike train, this ensures a constant number of non-zero-padded comparisons (*n_c_*) at each lag, as opposed to traditional cross-correlation in which *n_c_* is a function of lag. We divided the cross-correlogram for the segment by the mean value from 0.5–4 seconds on both sides, which allows the correlation’s units to be interpreted as the fraction of spikes greater than chance at a given lag (where 1 = chance). We repeated this process on overlapping segments (time step = 4 seconds) and then averaged these results together to get the mean, nonstationarity-corrected cross-correlogram. We generated a 99% confidence interval from the data with lag ≥ 0.5 s (which is a reasonable null distribution due to *n_c_*, and thus the variance of the correlation estimate, being held constant). We conclude that a pair is synchronous if its normalized cross-correlation at lag zero is larger than the upper boundary of this confidence interval.

### Behavioral Testing and Metric

Full details on the behavioral testing and the principal component analysis (PCA) metric for gradually depleted animals can be found in Willard et al. 2019. In brief, PCA was performed on the following metrics from behavioral tests: mean speed in an open field, number of rears in 10 minutes in a small enclosure, total time spent traversing a pole task, and latency to fall on a wire hang task.

### Linear Regression

Linear regression was performed using ordinary least squares. To determine if a linear fit was statistically significant, we computed 1000 fits each using a random subsample containing 80% of the data. We computed a bootstrapped confidence interval of the slope of this linear relationship from the middle 99% of the slopes of these 1000 fits, and the relationship was considered significant if this interval did not include zero.

### Decision Tree Regression

We sought to determine the relationship between dopamine loss, motor symptoms and neural firing by predicting animals’ TH immunofluorescence (see *Histology*) and the first principal component (PC1) of their behavior (see *Behavioral Testing and Metric*) from four physiological measures (see *Neural Measures*) and prevalence of delta oscillations. Firing rate, CV and bursts/second were averaged across all neurons for each animal, synchrony was measured as the fraction of synchronous pairs of units, and oscillations were measured as the fraction of delta oscillating units. Because of the highly nonlinear nature of these parameters’ relationships to dopamine loss and behavior (Willard et al., 2019), we used a variant of decision tree regression, a highly nonlinear regression method.

We built an individual tree on 80% of the data (20 animals) using the *fit* method of the DecisionTreeRegressor class in the scikit-learn package for Python to predict the percent of TH remaining (Y) from the above neuronal parameters (a set X). In brief, this method places all training data at the topmost node of a tree and calculates the mean squared error (MSE) of this node as if each animal’s TH were estimated to be the mean TH of every animal at the node. We determined, for each parameter X, the threshold T that would most reduce the mean squared error (MSE) of the animals if they were to be estimated in two different sets depending on whether their value of X is “greater than” or “less than or equal to” T. We then found the parameter for which the best T most reduces that MSE and split the animals at that node into two new child nodes according to the identified threshold. We iteratively repeated this process at every node until all terminal nodes had two or fewer animals at them, at which point each terminal node is termed a “leaf” of the tree.

We tested the remaining 20% of the data (5 animals) using the DecisionTreeRegressor *test* method, which runs each animal through the tree (picking > or ≤ at each node as determined by the animal’s data) until it reaches a leaf. The mean value of Y at each leaf is the prediction for that animal. We computed the error of the tree as the root-mean-squared error (RMSE) of its 5 predictions.

We computed a forest of 1000 such trees through subsampling the data into training and testing sets (Monte Carlo cross-validation) and calculated the top and bottom 2.5 percentiles to approximate a 95% confidence interval for the forest. We generated an intercept-only forest (using no parameters in the training set) and oscillation-only forest (using only the fraction of oscillations and an intercept term in the training set) on the same 1000 bootstrapped training and testing sets.

The importance of each parameter was determined using a variant on permutation importance. For a given parameter and tree, consider the set S of values for that parameter in the test set. We produced pseudo-test data with every derangement of S (i.e. 5 animals × 44 derangements of 5 values = 220 pseudo-test animals with shuffled data for one parameter). The difference between the RMSE of the real test data and the pseudo-test data is the importance of that parameter for that tree. To determine the parameter importance for the entire forest, we approximate a 95% confidence intervals as above from the 1000 trees.

A forest predicting the first principal component (PC1) of behavior instead of % TH remaining was computed in the same manner.

### ECoG-Spike Time Series Regression

To determine if SNr neurons have a significant lead/lag relationship with M1, we built a series of regression models predicting an M1 ECoG signal from the spiking of a single SNr unit at various lags. First, we binned the ECoG into 10ms bins and defined the dependent variable Y as the difference between adjacent ECoG measurements to reduce nonstationarity. We then built a 10th order autoregressive model of Y which served as the null model.

To incorporate SNr firing into the prediction, we calculated the spike density function (SDF) for an SNr unit by convolving its spike train with a Gaussian function with a standard deviation of 100 ms. We then aimed to determine which time shift of the SDF best improves the prediction of the ECoG. One might use a distributed lag model for this task, where the explanatory variables consist of the time shifted ECoG (autoregression) and all considered time shifts of the SNr SDF simultaneously in a single model, but the multicollinearity of the SDF at different time shifts can heavily bias the regression coefficients. Instead we assumed that, if a lag exists by which the unit firing influences the ECoG or vice versa, then there is only one such lag by which this influence occurs. Thus, we could build an individual model for each time shift of the SDF. Each model used the 10^th^ order autoregressive terms and one SDF term shifted from between −100 and +100 bins (−1000 to +1000 ms) as its explanatory variables. We built 201 such models, which covers the entire range of lags at 1 bin increments.

To determine if a significant lead/lag existed, we found the best model as determined by its mean squared error (MSE). We then determined if the model at this lag was significantly better than the null autoregressive model by performing an F-test at α < 0.05, correcting for 201 comparisons (Bonferroni correction). As choosing ECoG as the independent variable and using autoregressive terms from the past could introduce bias in favor of SNr predicting M1, we also performed these analyses using SNr as the independent variable (i.e. computing a null autoregressive model for SNr spiking and then computing 201 models at distinct ECoG time shifts to compare to the null), and performed the same analysis as above but in backwards time (i.e. building an autoregressive model of the ECoG from future ECoG samples). These analyses gave very similar results to the original analysis but were omitted for brevity.

### Statistical Tests

Statistical tests were performed to establish if fractions of oscillatory units and fractional ECoG bandpowers were significantly different across conditions. For comparisons with two groups, a two-sample t-test was performed, unless data were paired before and after a manipulation (e.g. acute drug infusion), in which case a one-sample t-test was performed. For comparisons with multiple groups compared against a control group, a one-way ANOVA was performed, and if this reached significance at the α = 0.05 level, a Dunnett’s post-hoc test was performed to determine if there were individual differences comparing groups to control. Asterisks above comparisons in figures correspond to *: p < 0.05, **: p < 0.01, *** p < 0.0001.

## Acknowledgements

Thanks to Rachel Bouchard, Hyun Young Park, Jenna Schwenk, Christen Snyder and Robert S. Turner for their help with this project. This work was supported by NSF awards DMS 1516288 (AHG, JER), 1724240 (JER), and NIH awards R01NS101016, R01NS104835, and R21NS095103 (AHG), and F31NS101821 (TCW)

---

**Figure 3 – Figure Supplement 1:**
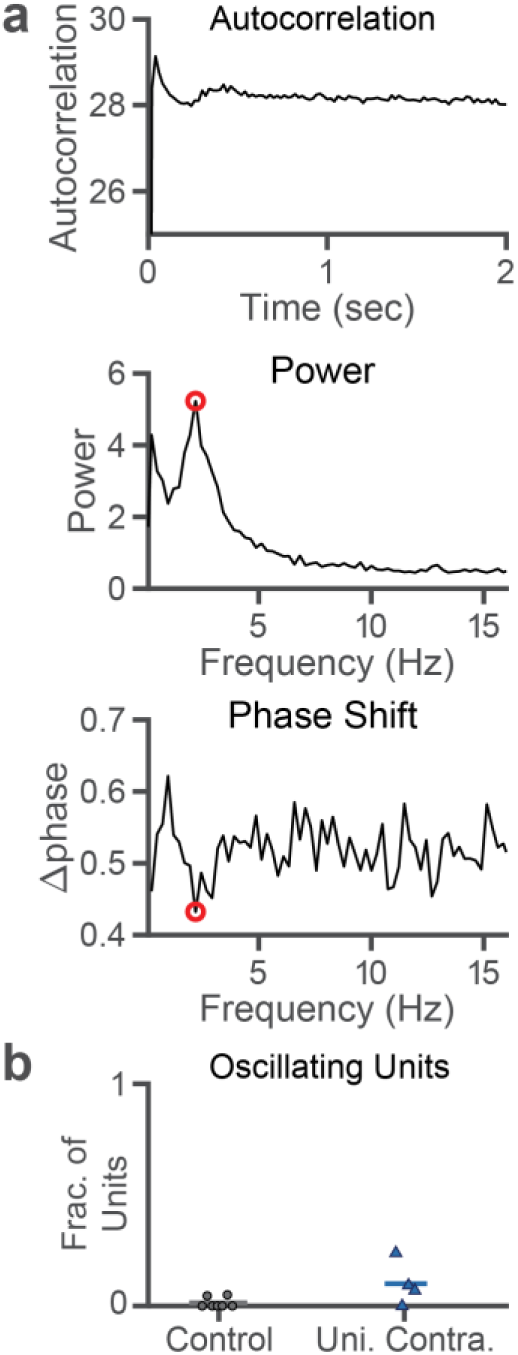
Unilaterally depleted animals exhibit a small number of delta oscillating units in the SNr of their dopamine intact hemisphere. **a.** Example autocorrelation (top), PSD (middle) and phase shift (bottom) for an example SNr unit exhibiting a delta oscillation in the intact hemisphere of a unilaterally depleted animal. **b.** Fraction of oscillating units in SNr for each control animal (black circle, n = 7) and in the intact hemisphere of unilaterally depleted animals (dark blue triangle, n = 4). The difference between these conditions is not significant at the α = 0.05 level (p = 0.1138, two-sample t-test).

**Figure 7 – Figure Supplement 1:**
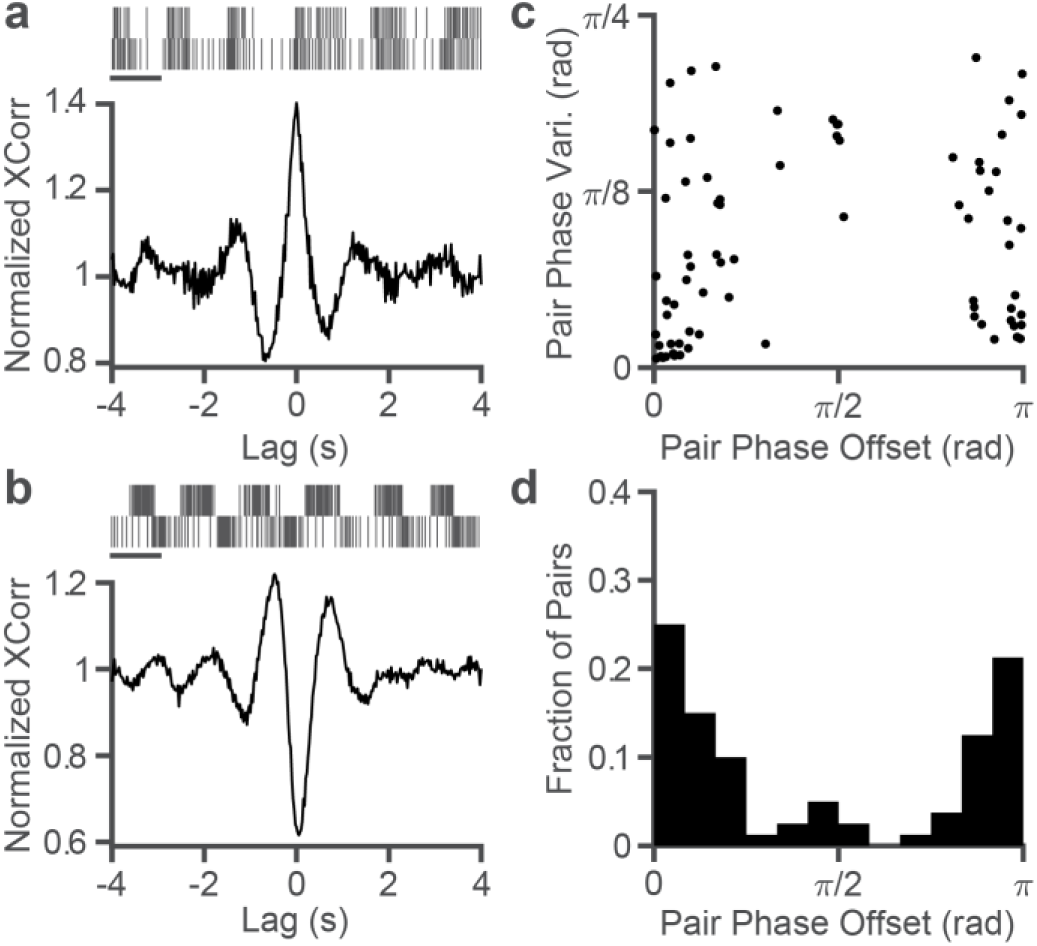
Pairwise phase relationships corroborate the existence of two populations of oscillating units in dopamine depleted SNr. **a.** Top: Spike rasters from a pair of simultaneously recorded SNr units, scale bar = 1 s. Bottom: Normalized cross correlations (see Neural Measures section of Methods) of the above pairs demonstrating an in-phase relationship. **b.** Same as **a** for a near anti-phase relationship. **c.** Scatterplot of all pairs of oscillating units. The horizontal axis measures their mean phase offset (0 indicating in phase, π indicating antiphase), and the vertical axis measures circular variance of phase offset computed across time windows. **d.** Histogram collapsing the above scatterplot to show counts of pairs based on their phase difference.

## Footnotes

1 Note that a rectangular window is used throughout this section. This is because 1) compared to tapered windows, the rectangular window’s maximal frequency resolution ensures a peak in the PSD representing an oscillation of interest will not mix with nearby peaks caused by pink noise, and 2) multiplication with a window function manifests as a convolution in the frequency domain, which distorts phase and yields a nearly flat and uninformative phase shift plot.

